# Long-term hematopoietic stem cells trigger quiescence in *Leishmania* parasites

**DOI:** 10.1101/2023.08.23.554403

**Authors:** Laura Dirkx, Sara Van Acker, Yasmine Nicolaes, João Luís Reis Cunha, Rokaya Ahmad, Ben Caljon, Hideo Imamura, Didier G. Ebo, Daniel C. Jeffares, Yann G.-J. Sterckx, Louis Maes, Sarah Hendrickx, Guy Caljon

## Abstract

Quiescence and posttreatment relapse constitute an important therapeutic constraint across the microbiological spectrum. This study unveils that *Leishmania infantum* and *L. donovani* parasites rapidly enter quiescence after an estimated 4-6 divisions in both mouse and human stem cells of the bone marrow but not in macrophages as primary host cells. Quiescent amastigotes display a reduced size and evidence for a rapid evolutionary adaptation response with genetic alterations. We formally demonstrate that acquisition of a quiescent phenotype endows parasites with a capacity to survive antileishmanial treatment. Transition through quiescence also results in an increased cellular infectivity and high transmission capacity through the sand fly vector. Transcriptional profiling of quiescent and non-quiescent metabolic states identified a limited set of 26 upregulated genes that are of particular interest given their predicted involvement as regulators of cell cycle progression and of gene expression at various levels. The differential gene set constitutes a reliable source for the identification of novel markers and potential drivers of quiescence, a metabolic state bestowing parasites the capacity to escape drug treatment.

## INTRODUCTION

Visceral leishmaniasis (VL) is a lethal neglected tropical disease caused by the obligate intracellular protozoan *Leishmania* [1, 2] and transmitted through the bites of infected female phlebotomine sand flies [3, 4]. *Leishmania* parasites alternate between two main morphological forms during their life cycle: a long flagellated extracellular promastigote within the sand fly and a non-flagellated obligate intracellular amastigote in the vertebrate host that resides within the monocyte-derived cells of the liver, spleen and bone marrow (BM) and eventually causes life-threatening complications [5–8].

The current antileishmanial drugs have many disadvantages and post-treatment relapse rates are increasing [9]. In many instances, relapse does not relate to reinfection, drug quality, drug exposure or resistance [10], but is rather due to persistence for which mechanistic information is lacking. Persistent infections can occur in a variety of host sanctuary tissues or cellular niches, such as hepatocytes (*Plasmodium vivax*), skeletal muscle and neurons (*Toxoplasma gondii*), adipose tissue (*Trypanosoma brucei* and *T. cruzi*) and the BM (*Mycobacterium tuberculosis*) [11–15]. Long-term hematopoietic stem cells (LT-HSC) in the BM were identified as a relapse niche for VL infection. LT-HSC become readily infected with extreme parasite burdens accompanied with low reactive oxygen species (ROS) and nitric oxide (NO) levels and a specific Stemleish transcriptional profile [16].

Besides persistence linked to cellular niches, treatment failure can also be associated with the adaptive behavior of parasites. In response to stress, certain microorganisms employ so-called quiescence to increase their chances of survival [17]. The quiescent state is characterized by a lowered metabolic activity and renders a microorganism tolerant to antibiotics at the expense of becoming non-proliferative [18, 19]. Hence quiescent cells are phenotypic variants of the wildtype, and their dormancy can be reversed when stressors are alleviated. Given its discernible clinical implications, microbial quiescence has gained considerable interest and has been the subject of intense research for certain pathogens, especially bacteria [18–22]. In contrast, quiescence in *Leishmania* has only recently been discovered and its role in drug tolerance, infection relapse and general parasite biology remains poorly understood. A recent study demonstrated that *Leishmania* quiescence can be induced by various triggers (*e.g.* antimonial drug pressure or stationary phase growth [23]). Furthermore, transcriptomic and metabolomic analyses of quiescent stages corroborated an overall downregulation of biosynthetic processes as a hallmark of quiescence [23]. Despite these important insights, the molecular determinants orchestrating the phenotypic transition to the quiescent state in *Leishmania* remain thus far unknown.

The present study started with the observation that visceral *Leishmania* amastigotes inside LT-HSC rapidly enter a quiescent state. Although the induction is unrelated to drug pressure, we demonstrate that these quiescent parasites benefit from an enhanced survival of antileishmanial treatment. To better understand the molecular basis underlying amastigote quiescence, we performed an unbiased total RNAseq, which led to the identification of genes that are significantly overexpressed in the LT-HSC-induced quiescent state. Functional annotation of the proteins encoded by these upregulated genes through AlphaFold2 structure prediction [24, 25] provide clues on the players involved in attaining the quiescent phenotype in a natural infection model. In addition, we show that transitioning through a quiescent state has a profound impact on parasite infectivity and transmissibility. The results provide mechanistic information on the *in situ* acquisition of quiescence and its downstream effects on parasite biology (survival under drug pressure, infection, and transmissibility). Given its broad relevance across the microbial spectrum, several of the quiescence-associated genes warrant future exploration as positive markers of quiescence/relapse or as putative targets in the transition process.

## RESULTS

### 1) *Leishmania* infection of mouse and human stem cells triggers amastigote quiescence

Serendipitously we discovered by flow cytometry that already after 24 hours of *Leishmania* infection in LT-HSC, there is a presence of two distinct DsRed^+^ amastigote populations (DsRed^hi^ and DsRed^lo^) suggesting different metabolic states (**Figure 1a**). The decreased DsRed signal indicates reduced expression from the 18S rDNA locus, previously reported as an indicator of entry into a quiescent state [26]. *In situ* amastigote quiescence was shown to occur independently of strain (*L. infantum* LEM3323 or clinical isolate LLM2346) or species (*L. infantum* and *L. donovani*) (**Figure 1a**) and was recorded in both mouse LT-HSC and human HSPC (**Figure 1b**). In contrast, amastigotes purified from infected macrophages cluster in one homogenous DsRed^hi^ population (**Figure 1c**). Promastigote back-transformation was used to confirm viability of sorted DsRed^lo^ parasites, the capacity to regain proliferative capacity and stability of the DsRed^lo^ phenotype. DsRed expression in the quiescent state remained lowered after transformation into the promastigote form (**Figure 1d**), which was also confirmed for derived monoclonal lines. Quiescent parasite cultures lost DsRed-expression after promastigote back-transformation with a frequency between 1.96% (1/51 clones) and 4.76% (2/42 clones) (**supplementary Figure 1**), suggesting that parasites undergo a rapid evolutionary adaptation response and genetic rearrangements upon entry and exit from quiescence. Loss of the dsRed gene was demonstrated in by PCR. In contrast, no clones derived from DsRed^hi^ parasites lost the DsRed construct. Interestingly, quiescent amastigotes exhibit a significantly reduced size compared to DsRed^hi^ amastigotes (**Figure 1e**). Confocal fluorescence microscopy confirmed heterogenous DsRed signals and variable amastigote size in the LT-HSC (**Figure 1f**). The mean fluorescence of the DsRed signal increased with the apparent amastigote size (forward scatter – FSC, **Figure 1g**), illustrating that acquisition of quiescence is associated with several cellular changes.

**Figure 1.**
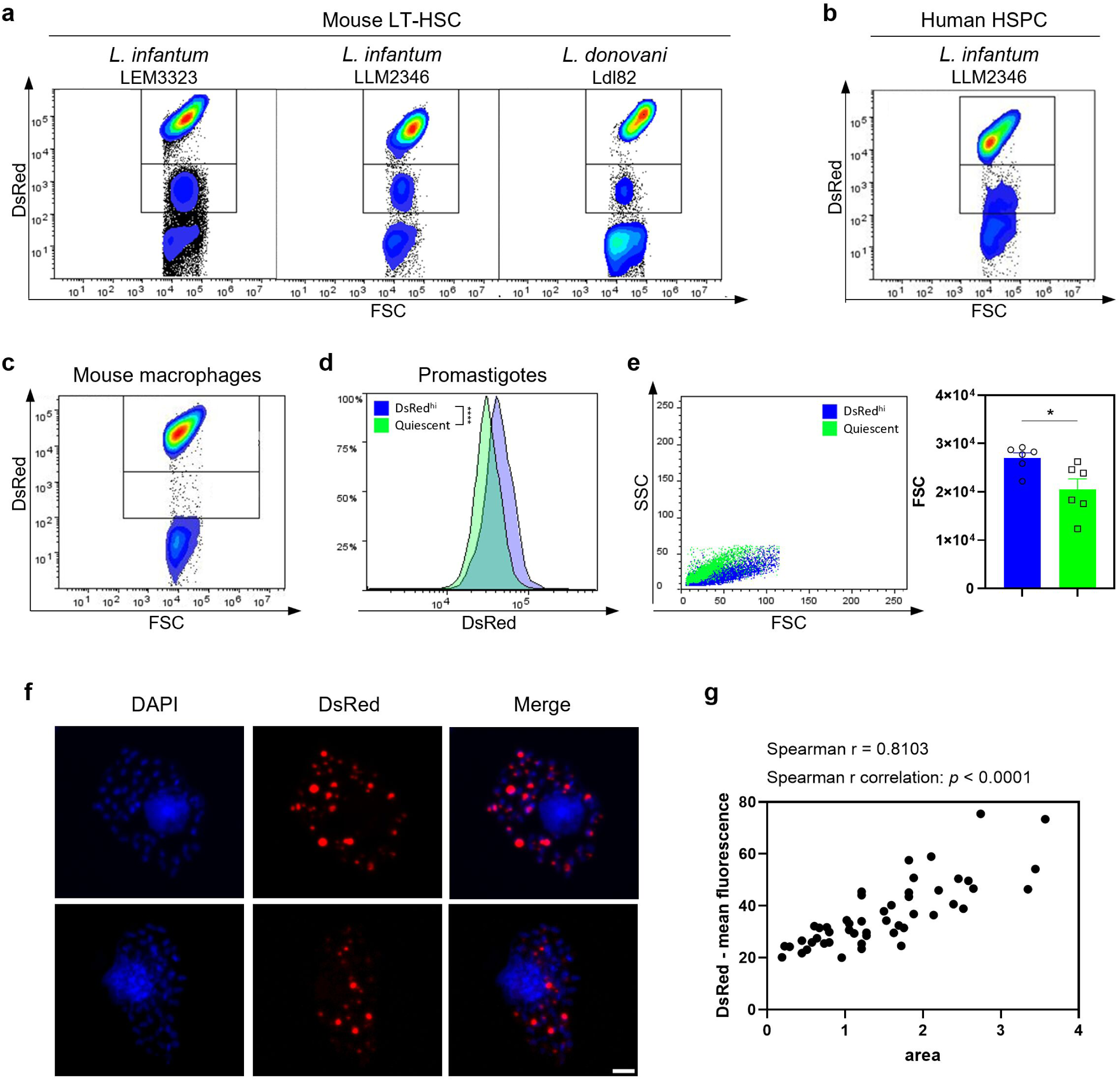
*Leishmania* infection of mouse LT-HSC and human HSPC triggers amastigote quiescence. **(a)** Amastigotes recovered from infected mouse LT-HSC and measured via flow cytometry. Cells in the left panel were infected with *L. infantum* LEM3323 WT^PpyRE9/DsRed^, middle panel with *L. infantum* clinical isolate LLM2346 WT^PpyRE9/DsRed^, right panel with *L. donovani* Ldl82 WT^PpyRE9/DsRed^. **(b)** Amastigotes recovered from *L. infantum* LLM2346 WT^PpyRE9/DsRed^ infected human HSPC. **(c)** BM derived macrophages were infected with *L. infantum* (LEM3323 WT^PpyRE9/DsRed^) and intracellular amastigotes were isolated and measured by flow cytometry. **(d)** DsRed expression measured by flow cytometry after promastigote back-transformation of DsRed^hi^ and DsRed^lo^ (*i.e.* quiescent) amastigotes recovered from LT-HSC. **(e)** *L. infantum* LEM3323 WT^PpyRE9/DsRed^ amastigotes recovered from infected mouse LT-HSC and measured via flow cytometry, back-gated on SSC versus FSC. Mann-Whitney test, \**p* < 0.05, six independent repeats. **(f)** Sorted mouse LT-HSC were infected with *L. infantum* (LEM3323 WT^PpyRE9/DsRed^) and processed for microscopy. DAPI (blue), amastigotes (red). Scale bar = 10 µm. **(g)** Analysis of microscopy images of (f) on ImageJ, comparing the expression level of DsRed to its respective size.

### 2) Amastigotes enter quiescence following *in situ* proliferation in mouse and human stem cells

To uncover why LT-HSC trigger the rapid development of quiescent amastigotes, a CFSE labelling of *L. infantum* LEM3323 promastigotes was performed to assess the number of parasite divisions before entering quiescence (*i.e.* acquire a DsRed^lo^ phenotype). The number of divisions was calculated based on a 2-fold decline at each division of the CFSE median fluorescence intensity (MFI) (**Figure 2a, left panel**). After 6 hours of co-incubation, the DsRed^hi^ amastigote fraction divided about 2 times compared to the control (0 hours), which was comparable to promastigote proliferation in 6-hour *in vitro* cultures. In contrast, the amastigote fraction that would eventually acquire a DsRed^lo^ phenotype displayed a more diverse pattern, ranging between 2, 4 and 6 *in situ* divisions (**Figure 2a)**. The highest proportion of amastigotes underwent 6 divisions to enter into quiescence (**Figure 2b**). These data indicate that the high proliferation rate in LT-HSC is a prevailing trigger. To confirm clinical relevance of these findings, amastigote quiescence was tested in human HSPC using a recent *L. infantum* clinical isolate (LLM2346 WT^PpyRE9/DsRed^). In **Figure 2c-d**, CFSE labelling of promastigotes was again performed to assess the number of divisions. After 24 hours of co-incubation, the DsRed^hi^ fraction divided about 0.5-1 time compared to the control (0 hours), which was comparable to the promastigote culture after 24 hours. The amastigote fraction that ultimately becomes quiescent divided about 4-5 times (**Figure 2d**). The observed difference in division rate between LEM3323 and LLM2346 can be linked to intrinsic growth rate differences (**Supplementary figure 2a, b**). Collectively these data show that *Leishmania* quiescence arises mostly after 4-6 divisions in mouse and human stem cells.

**Figure 2.**
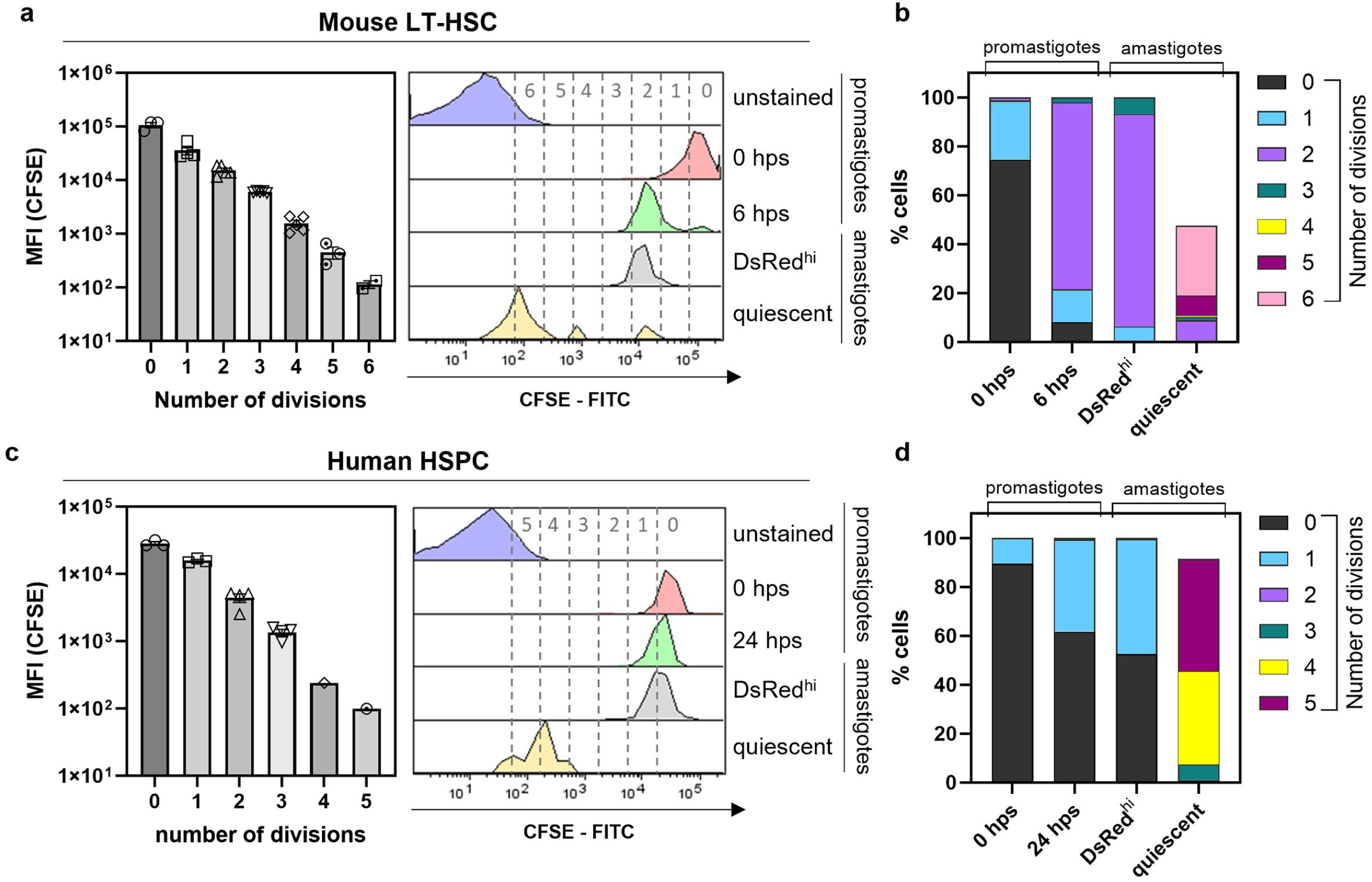
Number of divisions to trigger quiescence in amastigotes from mouse LT-HSC and human HSPC. (**a**) Number of divisions as calculated by CFSE staining and defined by halving of CFSE MFI (left panel). Controls are unstained and CFSE *L. infantum* (LEM3323 WT^PpyRE9/DsRed^) promastigote cultures. Sorted mouse LT-HSC were infected with CFSE labelled *L. infantum* promastigotes and amastigotes were recovered after 6 hours of co-incubation (right panel). **(b)** Percentage of cells in each division range from (a). Results are based on three independent repeats. **(c)** Number of divisions as calculated by CFSE staining and defined by halving of CFSE MFI (left panel). Controls are unstained and CFSE *L. infantum* (LLM2346 WT^PpyRE9/DsRed^) promastigote cultures. Sorted human HSPC were infected with CFSE labelled *L. infantum* promastigotes and amastigotes were recovered after 24 hours of co-incubation (right panel). **(d)** Percentage of cells in each division range from (c). Results are based on three independent repeats.

### 3) Transition through *in situ* quiescence enhances parasite survival, infectivity, and transmission potential

Next, we wondered whether adoption of a quiescent state would have an impact on various aspects of parasite biology. To investigate whether amastigote quiescence in LT-HSC is linked to survival of treatment and consequent occurrence of relapse, sorted/infected cells were treated with 250 µM paromomycin (PMM) or 7.5 µM miltefosine (MIL) for 72 hours before purifying the remaining amastigotes and determining the distribution of quiescent parasites based on DsRed fluorescence by flow cytometry. Drug treatment was found to primarily affect DsRed^hi^ parasites, increasing the proportion of quiescent amastigotes (**Figure 3a,b**).

**Figure 3.**
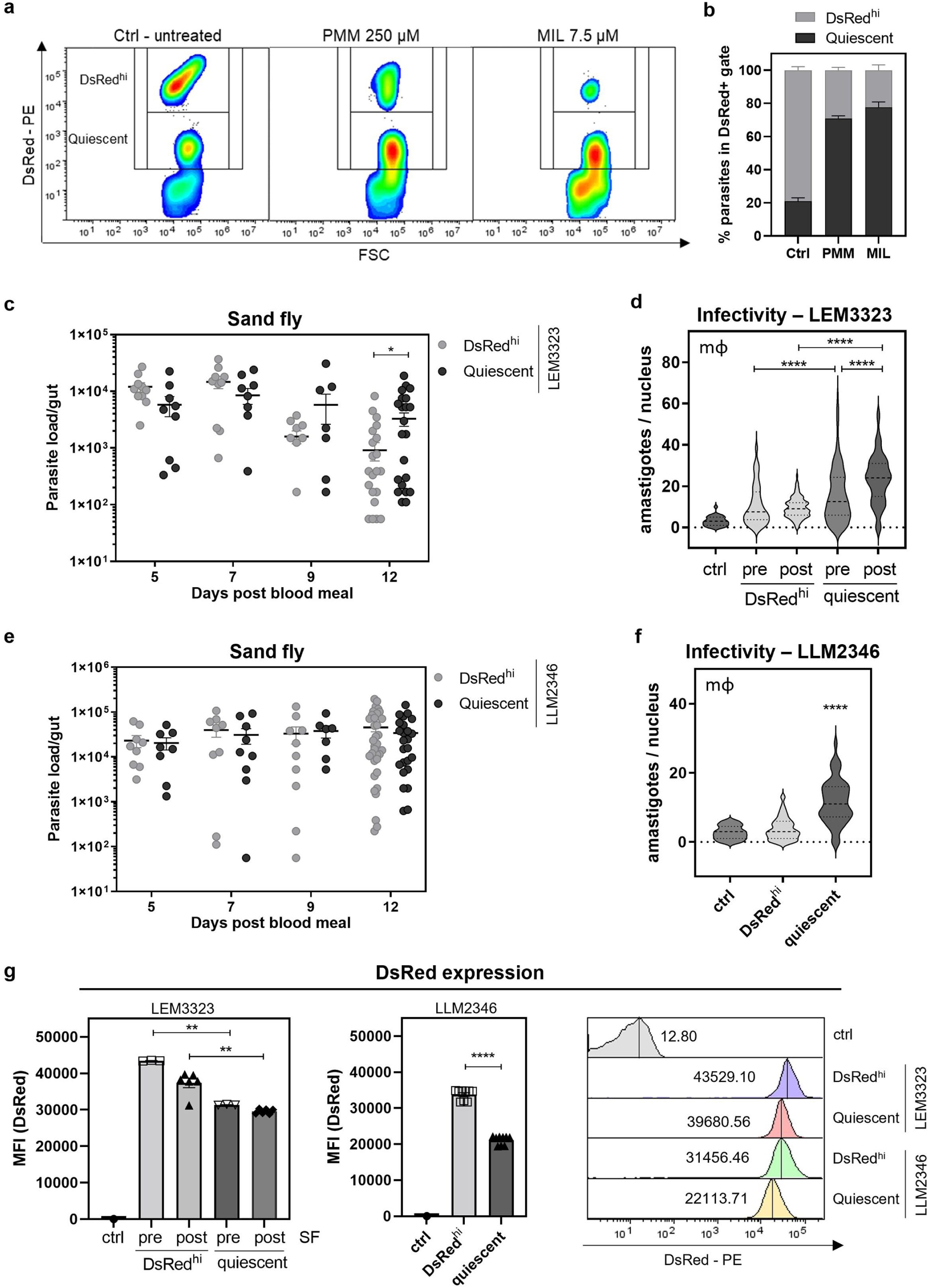
Phenotypic characteristics of quiescent amastigotes from LT-HSC. (**a-b**) Sorted LT-HSC were infected with *L. infantum* (LEM3323 WT^PpyRE9/DsRed^) for 24 hours followed by treatment with 250 µM PMM or 7.5 µM MIL for 72 hours. To compare pre– and post-treatment distribution of quiescent parasites, amastigotes were isolated and remeasured on the FACSMelody. **(c)** Sand flies were infected by LEM3323 promastigotes recovered from DsRed^hi^ and quiescent amastigotes in mouse LT-HSC. The parasite load in the gut was assessed at day 5, 7, 9 and 12 after infection (blood meal). Sand fly infections were repeated three independent times. Unpaired *t* test, 10 < *n* < 30, \**p* <0.05. **(d)** LEM3323 promastigotes of a control culture and promastigotes recovered from DsRed^hi^ and quiescent parasites in mouse LT-HSC, pre and post sand fly passage (SF) were co-incubated with peritoneal macrophages for 96 hours and infectivity was assessed with Giemsa staining. The original promastigote culture was used as control and was significantly lower than pre SF DsRed^hi^ (*p* <0.01), post SF DsRed^hi^ (*p* <0.05), pre and post SF quiescent (*p* <0.0001). Ordinary one-way ANOVA, 30 < *n* < 130, \*\*\*\**p* <0.0001. **(e)** Sand flies were infected by LLM2346 promastigotes recovered from DsRed^hi^ and quiescent amastigotes in human HSPC. The parasite load in the gut was assessed at day 5, 7, 9 and 12 after the infectious blood meal. Sand fly infections were repeated three independent times. 10 < *n* < 34. **(f)** LLM2346 promastigotes of DsRed^hi^ and quiescent recovered promastigotes from human HSPC were co-incubated with peritoneal macrophages for 96 hours and infectivity was assessed with Giemsa. Mann-Whitney test, *n* = 100, \*\*\*\**p* <0.0001. **(g)** DsRed expression of LEM3323 DsRed^hi^ and quiescent promastigotes recovered from mouse LT-HSC before or after passage through the sand fly, and LLM2346 DsRed^hi^ and quiescent promastigotes recovered from human HSPC. Mann-Whitney test, \*\**p* <0.01. All experiments are expressed as mean ± SEM.

To assess whether going through a reversible quiescent state influences subsequent transmission, sand flies were infected with promastigotes derived from DsRed^hi^ and quiescent amastigotes. In **Figure 3c**, parasite load in the gut was compared at different time points, with parasites having transitioned through a quiescent state showing a slightly enhanced sand fly infectivity (*p* <0.05 at day 12). A similar analysis was performed for parasites derived from human HSPC, where infectivity in the sand fly vector remained the same (**Figure 3e**).

As shown before sand fly passage, quiescent parasites post sand fly showed a lower DsRed signal (**Figure 3g**), indicating that some quiescence-associated phenotypic changes are stable after sand fly passage. Infectivity was evaluated by co-incubation with peritoneal macrophages for 96 hours. Interestingly, promastigotes derived from quiescent (DsRed^lo^) amastigotes had a significantly higher infectivity compared to their DsRed^hi^ counterparts (*p* <0.0001) or the original promastigote culture before LT-HSC passage (*p* <0.0001). This difference in infectivity was even more pronounced after sand fly passage (**Figure 3d**). The infectivity in macrophages was also observed to be significantly higher for promastigotes derived from the quiescent strain recovered from human HSPC (*p* <0.0001) than for DsRed^hi^ promastigotes (**Figure 3f**). The *in vitro* growth curves of LEM3323 and LLM2346 promastigotes comparing quiescent and DsRed^hi^ phenotypes show no significant differences, suggesting that growth rate is not a determining factor contributing to the observed higher infectivity (**supplementary Figure 2**). These results highlight that a transition through quiescence not only affects treatment but also significantly influences important life cycle features such as infectivity and transmissibility.

### 4) *In vivo* relapse parasites share characteristics with parasites that transitioned through quiescence

Using a previously described reproducible post-treatment relapse model [16], mice were sacrificed at 6 weeks post infection (4 weeks post PMM treatment) and BM was collected for promastigote back-transformation (**Figure 4a**). Infection of mouse peritoneal macrophages revealed an enhanced infectivity of relapse parasites (**Figure 4b**) as was found for quiescent parasites. The percentage of reduction after *in vitro* PMM treatment of infected macrophages remained stable for relapse versus the parental parasites, demonstrating that these parasites did not acquire a drug resistant phenotype (**Figure 4c**). To check whether the parasites were still infective for sand flies, the gut parasite load was compared at different time points. No significant differences in sand fly parasite loads (**Figure 4d**) and *in vitro* promastigote growth (**Supplementary** Figure 2c) were detected. The recorded high fitness of relapse parasites overlaps with the parasite phenotype after transition through a quiescent stage in LT-HSC.

**Figure 4.**
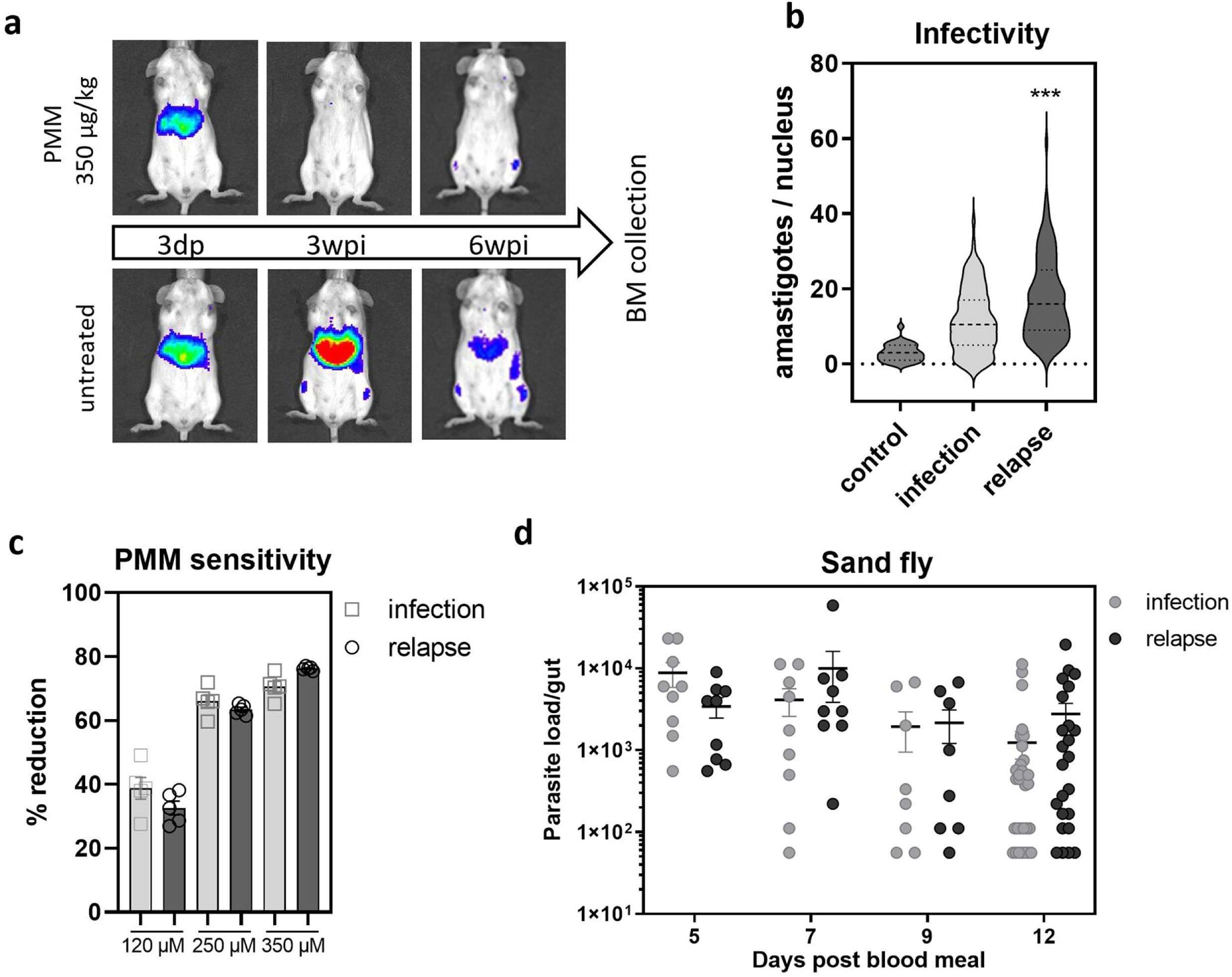
Phenotypic characteristics of promastigotes from relapsed BM. (**a**) BALB/c mice were infected with 10^8^ metacyclic promastigotes of *L. infantum* LEM3323 WT^PpyRE9/DsRed^. One group was treated with PMM 350 µg/kg per day (IP) for 5 consecutive days. Infection was followed up by BLI, BM was collected at 6 weeks post infection (wpi) from untreated and relapsed mice. **(b)** Promastigotes recovered from relapsed BM (relapse) and untreated BM (infection) were co-incubated with peritoneal macrophages for 96 hours and infectivity was assessed with Giemsa. Mann-Whitney test, *n* = 100, \*\*\**p* <0.001. **(c)** Promastigotes recovered from relapsed BM (relapse) and untreated BM (infection) were co-incubated with peritoneal macrophages for 24 hours and treated with 120 µM, 250 µM or 350 µM PMM for 72 hours, infectivity was assessed with Giemsa. **(d)** Sand flies were infected by promastigotes recovered from relapsed BM (relapse) and untreated BM (infection) and the parasite load in the gut from infected flies was assessed at day 5, 7, 9 and 12 after infection (blood meal). Sand fly infections were repeated three independent times. 10 < *n* < 32.

### 5) *In situ* quiescent amastigotes undergo vast transcriptional changes and reveal new markers and potential drivers

To unravel the molecular basis for quiescence in LT-HSC amastigotes, unbiased total RNA sequencing was performed on three independent samples of 10,000 DsRed^lo^ (quiescent) and DsRed^hi^ (non-quiescent) amastigotes that were isolated and flow sorted from mouse LT-HSC in three independent infection experiments. Principal component analyses (**Figure 5a, b**) revealed distant profiles of the quiescent and DsRed^hi^ samples, supporting the observed difference between both parasite phenotypes. Consistent with a previous quiescence study [23], ribosomal genes were strongly downregulated. The ribosomal genes (194 genes) and transfer RNA (37 genes) were removed for the downstream differential expression analysis. Following trimming of the dataset, 1258 genes were found differentially expressed (*padj* <0.05) in quiescent amastigotes (24 up– and 1234 downregulated, **Figure 5c** and Supplementary Data xlsx-file). Beyond structural constituents of the ribosome, a large proportion of downregulated genes was enriched in biological processes such oxidation-reduction and various other metabolic processes (ATP, carbohydrate derivatives, nucleobase-containing small molecules, L-methionine and ethanolamine biosynthesis), mitochondrial transport, organization and electron transport, cellular protein localization, response to a temperature stimulus and motility (ciliary/flagellar basal body organization) (**Supplementary** Figure 3). The top 25 downregulated genes (5 hypothetical and 20 with annotation), from which 22 were overlapping in all three replicates, may serve as negative markers of quiescence (**Figure 5f**, **Table 1)**. The previously described absence of an oxidative response to *Leishmania* in the LT-HSC niche and the induction of quiescence may explain why oxidoreductase activity is downregulated [16].

**Figure 5.**
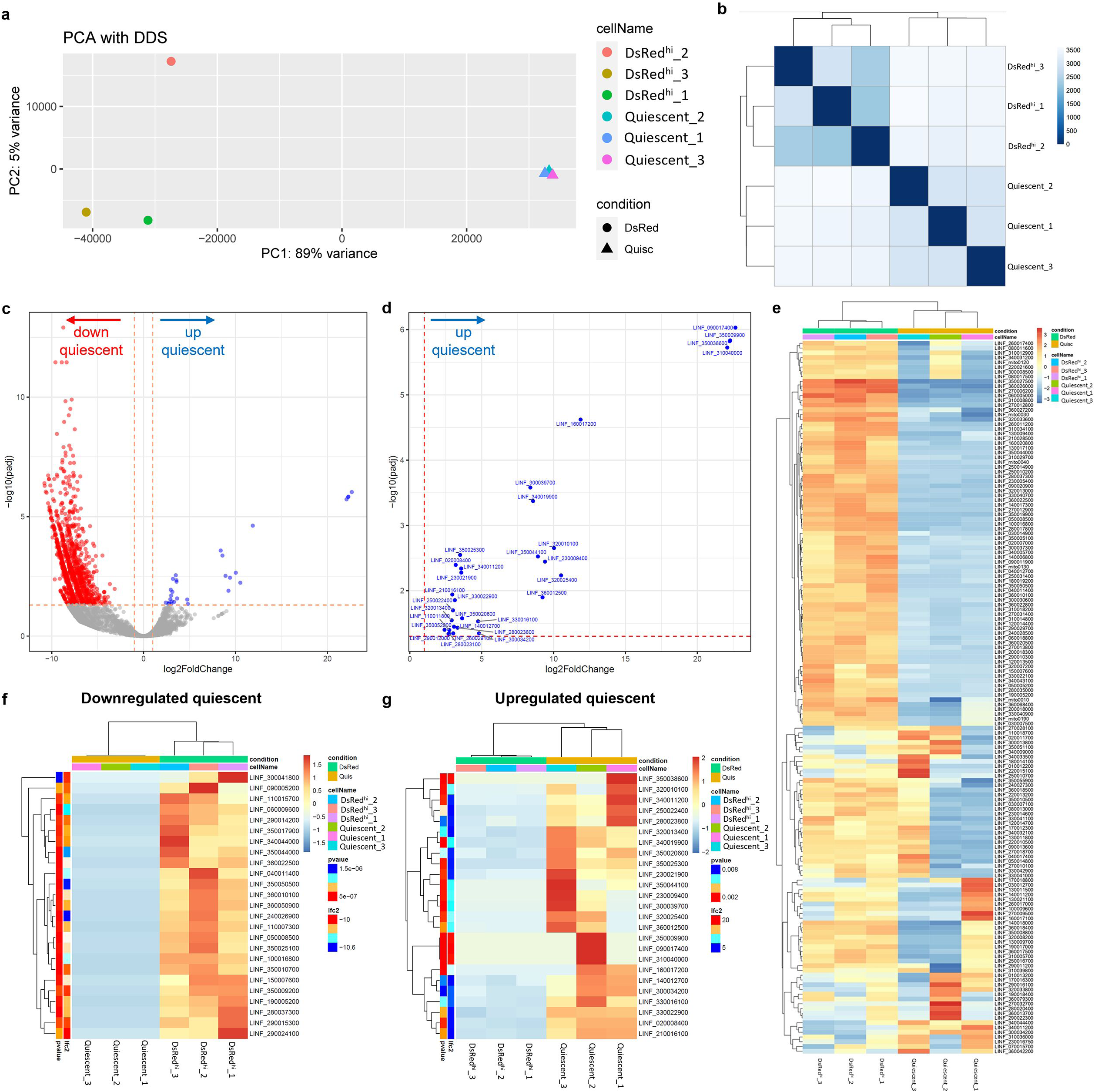
RNA seq data from quiescent amastigotes recovered from mouse LT-HSC. (**a**) Principal component analysis (PCA) of the RNA seq data revealing distant clustering of the independent DsRed^hi^ and quiescent samples. **(b)** Euclidean distance matrix between the samples illustrating the Poisson Distance. **(c)** Volcano plot from quiescent amastigotes displaying significantly up– and downregulated genes. **(d)** Zoom in the top-right section of (c), showing the gene IDS. **(e)** Heatmap of the top 150 differentially expressed genes. **(f)** Top 25 downregulated genes in quiescent amastigotes. **(g)** Top 25 upregulated genes in quiescent amastigotes. **(e-g)** Expression levels are represented with a color scale ranging from blue to red (overexpression). These heatmaps were generated excluding tRNA and transcripts encoding ribosomal proteins.

**Table 1.**
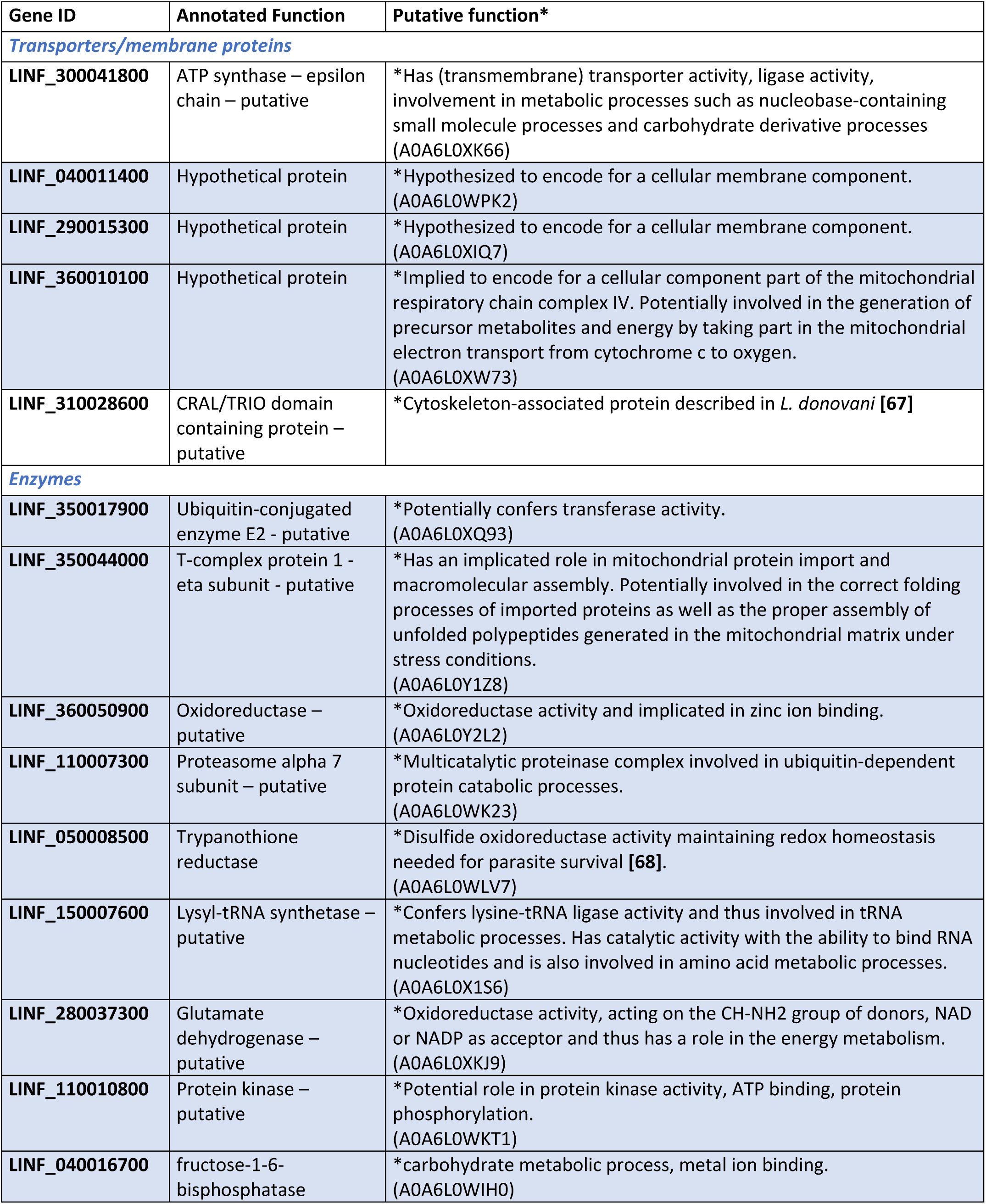
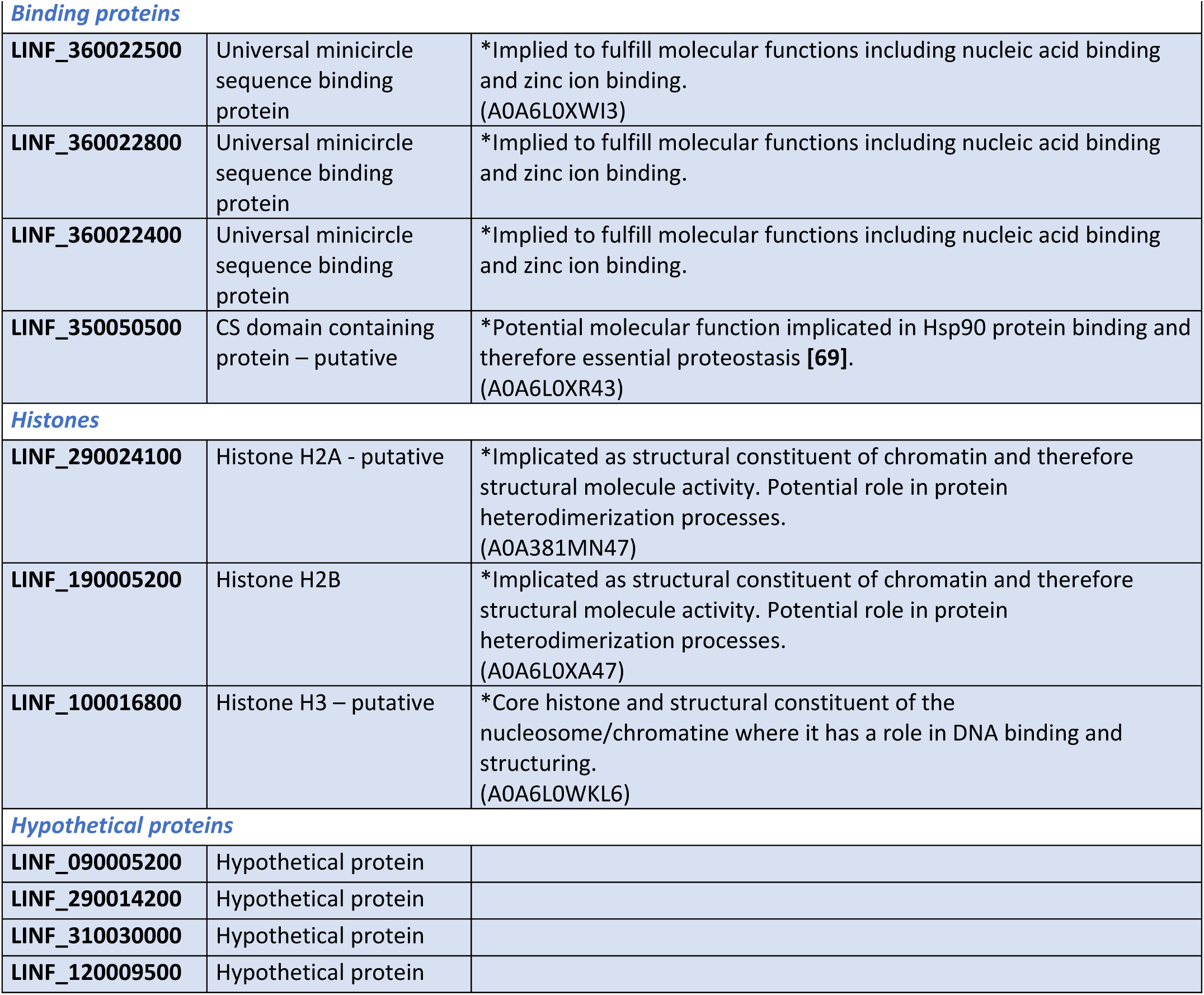
Top downregulated genes in quiescent amastigotes. Shaded cells represent congruent downregulation in the three independent biological replicates. Uniprot accession codes are indicated between brackets.

Interestingly, 24 genes in the quiescent replicates were significantly upregulated considering a log2-fold change and *padj* <0.05 (**Figure 5d**), of which 14 showed clear congruence across the three independent biological replicates (**Figure 5e, g**). From this list of 24 signature quiescence genes, only 9 have been functionally annotated, whereas the remaining 15 are labeled as “hypothetical”. We reverted to AlphaFold2 structure prediction followed by searches with the DALI server to gain insights into the biological functions of the proteins encoded by these upregulated quiescence-associated genes. Four out of 24 candidates (LINF_230009400, LINF_330016100, LINF_350009900, LINF_350020600) were omitted from further investigation due to poor overall pLDDT scores or the absence of a clear fold in the predicted structures. For those structure predictions characterized by good to excellent overall pLDDT scores, functional annotation was relatively straightforward. For others, functions are proposed based on specific protein regions/domains displaying a reasonable to excellent model quality (*e.g.* LINF_250022400 and LINF_350025300). The structure-based functional annotation for these proteins is summarized in **Figure 6** and **Table 2**. The analysis reveals that the proteins encoded by the upregulated quiescence-associated genes are involved in a myriad of cellular processes: metabolism, regulation of gene expression at the RNA level, cell motility and cytoskeleton dynamics, pleiotropic mediators of protein-protein interactions, vesicle transport and autophagosome, and DNA topology and cell cycle control. These results suggest that the phenotypic transition to and/or maintenance of quiescence requires a coordinated cell-wide response.

**Figure 6.**
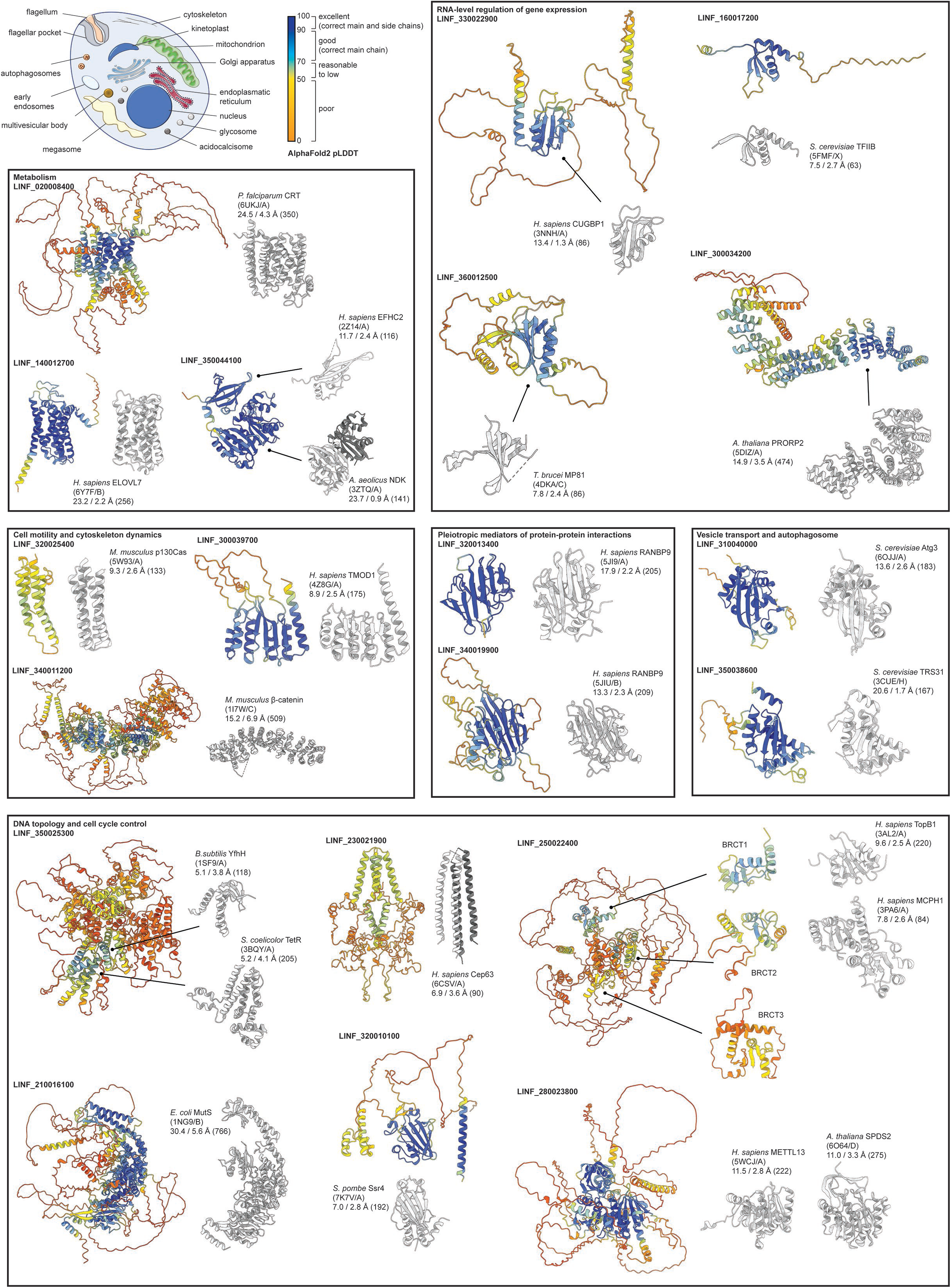
Functional annotation of upregulated quiescence-associated genes through AlphaFold2 structure prediction and structural homology searches. The top left corner depicts a cartoon of an amastigote. The boxes contain cartoon representations of the AlphaFold2 structure predictions, which have been grouped together based on their structure-based functional annotation (*cfr*. Table 2). The structures are colored according to the predicted local distance difference test (pLDDT) score, which reflects (local) model quality. The most representative DALI hits are shown in a cartoon representation and have been colored light grey. The accompanying text provides information on the nature of the DALI hit (first line), the PBD code and chain ID (second line), and the DALI Z-score, Cα rmsd and number of residues included in the structural overlap (third line).

**Table 2.** Top upregulated genes in quiescent amastigotes. Shaded cells represent congruent upregulation in the three independent biological replicates. Green shaded cells represent genes for which structural and functional predictions are reliable, blue shaded cells represent genes for which annotating one single function is difficult. Orange shaded cells represent predictions with low confidence.

| Gene ID | seq | Annotated Function | Structural homology and predicted function* |
| --- | --- | --- | --- |
| *DNA topology and cell cycle control* |  |  |  |
| LINF_210016100 | 4110 | Mis-match repair protein – putative | * DNA binding and mismatch repair |
| LINF_230021900 | 921 | Hypothetical protein | Centrosomin<br>*Role in regulating cell cycle progression and cell division |
| LINF_250022400 | 3507 | Hypothetical protein | DNA topoisomerase 2-binding protein, contains 3 BRCT domains (2 clearly predicted and 1 with poor accuracy) and in that regard resembles microcephalin (implicated in chromosome condensation and DNA damage induced cellular response).<br>Interpro accession code IPR001357<br>*DNA repair, recombination, topology and cell cycle control |
| LINF_280023800 | 2823 | Hypothetical protein | S-adenosyl-L-methionine-dependent methyltransferases/spermidine synthases/polyamine aminopropyltransferase.<br>Interpro accession code IPR029063<br>*Pleiotropic functions including cell cycle control |
| LINF_320010100 | 1152 | Hypothetical protein | SWI/SNF and RSC complexes subunit SSR4 / DNA binding (contains a DNA binding domain conserved in some transcription factors and a Zn<sup>2+</sup> coordination center)<br>*Chromatin remodeling, regulation of transcription |
| LINF_330016100 | 1026 | Hypothetical protein | * Regulation of gene expression |
| LINF_350009900 | 1017 | Hypothetical protein | Chromodomain-helicase-DNA-binding protein 1<br>* Chromatin remodeling |
| LINF_350025300 | 4260 | Hypothetical protein | Contains well-modelled regions with structural homology to DNA binding domains |
| *Pleiotropic mediators of protein-protein interactions* |  |  |  |
| LINF_320013400 | 498 | Hypothetical protein | SPRY domain containing protein |
| LINF_340019900 | 1041 | Hypothetical protein | SPRY domain containing protein |
| *RNA-level regulation of gene expression* |  |  |  |
| LINF_160017200 | 354 | Hypothetical protein | RNA polymerase II transcription factor B subunit<br>* Regulation of transcription |
| LINF_300034200 | 2022 | Pentatricopeptide repeat domain containing protein – putative | Large family of modular RNA-binding proteins in kinetoplasts which mediate several aspects of gene expression primarily in mitochondria but also in the nucleus<br>* RNA binding, RNA processing in mitochondria [70] |
| LINF_330022900 | 993 | RNA recognition motif – putative | * RNA binding, post-transcriptional regulation of gene expression (RNA recognition motif) |
| LINF_350020600 | 5577 | Hypothetical protein | GTP-binding nuclear protein RAN<br>Importing beta-like protein KAP120<br>* Nuclear translocation of RNA and proteins |
| LINF_360012500 | 807 | KREPA4 – putative | * RNA binding protein, catalyzing the editing of most mitochondrial mRNAs. KREP4A is essential for editosome integrity and survival of *Trypanosoma brucei* [71] |
| **Metabolism** |  |  |  |
| LINF_020008400 | 2919 | EamA-like transporter family/Triose-phosphate Transporter family/UAA transporter family – putative | Transmembrane transport - Multi-pass membrane protein<br>* Drug/metabolite transporter superfamily<br>* Putative increased tolerance to drugs |
| LINF_140012700 | 981 | Fatty acid elongase - putative | * Lipid metabolism, *Leishmania* polyunsaturated fatty acid metabolites are implicated in parasite/host interactions (macrophage M2 polarization) |
| LINF_350044100 | 1014 | Nucleoside diphosphate kinase - putative | NDP kinase (this gene product has two NDP domains and an EF-hand domain to allow dimerization).<br>* Nucleotide metabolism and host cell protection from extracellular ATP [72] |
| **Vesicle transport and autophagosome** |  |  |  |
| LINF_310040000 | 567 | Autophagy protein 10 (ATG10) | Structural homology to ATG3<br>* Autophagosome formation |
| LINF_350038600 | 630 | Transport protein particle (TRAPP) subunit - putative | * ER-to-Golgi transport of vesicles, adaptation of cycling cells to stress conditions [73] |
| LINF_230009400 | 465 | Hypothetical protein | * Delivery of tail-anchored (TA) proteins to the endoplasmic reticulum |
| **Cell motility and cytoskeleton dynamics** |  |  |  |
| LINF_340011200 | 1749 | Hypothetical protein | * Flagellar assembly and stability |
| LINF_300039700 | 705 | Hypothetical protein | Tropomodulin/leiomodulin/Actin<br>* Regulation cytoskeletal dynamics |
| LINF_320025400 | 351 | Hypothetical protein | Enhancer of filamentation 1 (focal adhesion domain; bacterial complement inhibitory domain)<br>* Disassembly of the primary cilium/flagellum by activation of Aurora A kinase at the basal body [74] |

## DISCUSSION

Current resources for new drug development for VL are scarce. Despite the imminent need to combat the increasing relapse rates, little is known on the underlying cause and suitable methods to study this phenomenon. Notably, post-treatment relapse is mostly not due to reinfection, drug quality, drug exposure or drug-resistant parasites [10, 27], but rather due to persistence of the pathogen. This persistence causing subclinical infections and subsequent relapse has been widely described across the microbiological spectrum [11–15, 28–30]. Generally, two aspects can be the cause of this: pathogens residing in sanctuary sites or niches, or pathogens switching to a quiescent phenotype. The first allows pathogens to survive and escape treatment or immunity without genetic or phenotypic changes. The latter relies on phenotypic diversity, *e.g.* quiescent or dormant forms [31]. Based on our observations, treatment failure in VL most likely results from an intrinsic ability of the *Leishmania* parasite to benefit from both quiescence and the occupation of sanctuary sites. We have previously identified a relapse-prone cellular niche in the BM in which treatment is less effective [16]. Here, we formally link quiescent parasites residing in this particular sanctuary site to drug tolerance.

In literature there is no clear consensus on the definition of persistence or quiescence, and additional synonyms such as dormancy and latency are used interchangeably as conceptually related terms. More commonly, quiescence refers to genetically drug susceptible, dormant (non-growing or slow growing) organisms that survive exposure to a given cidal drug and have the capacity to regrow (or resuscitate and grow) under highly specific conditions [32]. Several external triggers may induce quiescence, such as host immunity, drug pressure and/or nutritional and energetic stress [20, 33]. Investigation of the physiology of this phenotype *in vivo* is very challenging due to their scarcity and difficulty of detection. A review by Barrett *et al.* envisages several methods to study quiescence such as fluorescent probes, DNA replication probes, sorting of quiescent *vs* replicating cells, and several omics approaches [31]. Studies in various model organisms have been proposed to gain insights in the enigmatic basis of quiescence, such as transgenic fluorescent hypnozoites for *P. vivax* [34], multiple-stress model to study *M. tuberculosis* [35], and GFP reporter gene expression from an 18S rDNA locus in cutaneous leishmaniasis [26].

Quiescence in (cutaneous) leishmaniasis has been observed in several experimental systems since the first observation in 2015 by Kloehn *et al.* [36]. A form of quiescence has been described in axenic cultures of cutaneous *Leishmania* strains treated with antimonials or grown under stress conditions [23, 26]. A recent study on *L. mexicana* dermal granulomas describes a mosaic of metabolically active and semi-quiescent parasites during acute phases of infection, linking the phenotype to treatment failure [37]. Another study uses *in vivo* labeling with bromodeoxyuridine (BrdU) to visualize persistent slow-growing *L. major* parasites in macrophages in the skin [38]. To date, a knowledge gap exists for naturally elicited quiescent parasites recovered from a host cell niche, especially for VL. The present study provides unprecedented insights into quiescence of VL parasites in LT-HSC. The *Leishmania* amastigote stage had already been described as a less active state that may represent an adaptive response to a growth-restrictive intracellular microenvironment in granulomas [36, 39]. Here, we compared amastigotes within the common macrophage host cell to those within the LT-HSC niche and found distinct differences. Within LT-HSC, quiescence occurred within 6 hours following an estimated 4-6 divisions that affect amastigote size and are linked to rapid genetic alterations as observed by the erosion of the dsRed expression cassette. This indicates that host cell differences affecting the intracellular parasite proliferation rate determine the occurrence of quiescence.

Characterizing quiescent VL parasites recovered from LT-HSC showed an increased infectivity and a high capacity to colonize the sand fly gut. Consistent with the quiescent phenotype, parasites recovered from the BM of relapsed mice show an increased fitness with an elevated macrophage infectivity combined with a high sand fly infectivity. This suggests that the selected phenotype, typically associated with transition through a quiescent state, may pose an additional threat to leishmaniasis control programs. An increased infectivity associated with relapse of *L. donovani* infection has already been shown for miltefosine and antimonial treatment in the Indian subcontinent [10, 40, 41]. To prevent treatment failure, we advocate that the quiescent state of *Leishmania* should be also considered in the early stages of the drug discovery process.

Transcriptomic analysis of quiescent VL parasites revealed an overall downregulation of RNA, especially ribosomal protein genes, a signature of quiescence well described in literature [23, 36, 42, 43], as ribosome biosynthesis is one of the most energy intensive processes in the cell and thus a measure of the metabolic state. A large proportion of genes was downregulated in quiescent VL parasites, enriched in biological processes such oxidation-reduction and various other metabolic processes (ATP, carbohydrate derivatives, nucleobase-containing small molecules, L-methionine and ethanolamine biosynthesis), mitochondrial transport, organization and electron transport, cellular protein localization, response to a temperature stimulus and motility (ciliary/flagellar basal body organization). Metabolic changes in protozoan persisters are corroborated by other literature reports. Studies have shown that DNA replication, general transcription and protein synthesis are decreased in *Plasmodium spp*. and *Toxoplasma gondii* persisters [31]. The artificial axenic amastigote forms of *L. mexicana* and *L. braziliensis* showed downregulated synthesis of ATP, ribosomal components, proteins and alterations in membrane lipids [26, 39]. In contrast to many other species, *Leishmania* does not have transcriptional regulation and as such is mainly compensated by post-transcriptional mechanisms by RNA-binding proteins [44]. The observed downregulation in translation initiation factor activity, critical for gene expression, together with lower unfolded protein binding could explain the decreased DsRed expression observed in quiescent parasites. The overall downregulated cellular components are related to the energy metabolism (mitochondrion, mitochondrial part, mitochondrial membrane, respiratory chain complex IV). Mitochondrial gene expression undergoes changes according to the cell culture conditions and has been described to be reduced in an artificial model of quiescence [23].

Although quiescent forms exhibit an overall gene downregulation, some processes are sustained (*e.g.* oxidation-reduction in *T. gondii*, *M. tuberculosis* and *P. cynomolgi* hypnozoite forms [45–47]) or even upregulated (*e.g.* autophagy, amino acid catabolism, GP63 and amastin surface-like proteins in *L. braziliensis* axenic amastigote forms [23]). We also identified genes that were consistently upregulated in our *in situ* quiescence model and our structure-based functional annotation reveals that the encoded proteins are involved in a myriad of cellular processes. Several genes appear to be involved in DNA topology and cell cycle control (LINF_210016100, LINF_230021900 LINF_250022400, LINF_280023800, LINF_320010100, and LINF_350025300). The regulation of gene expression at the RNA level is also involved in acquiring and/or maintaining quiescence and differential transcript data also suggest alterations in RNA editing (LINF_160017200, LINF_300034200, LINF_330022900, LINF_360012500). As mentioned above, regulation of gene expression for kinetoplastids is by RNA level regulation and editing. Indeed, *Leishmania* is known for its extreme genomic and phenotypic variability, whereby a rapid shift in the gene repertoire is one of the key mechanisms for swift adaptation to changing environments [48], some of which have been associated with increased fitness in stress conditions or drug resistance [49]. Interestingly, some upregulated genes are involved in processes such as cell motility and cytoskeleton dynamics (LINF_300039700, LINF_320025400, and LINF_340011200), and vesicle transport and the autophagosome (LINF_310040000 and LINF_350038600). The latter is consistent with autophagy being upregulated in *L. braziliensis* axenic amastigote forms [23]. Two genes encode proteins with an SPRY domain, which is typically found in scaffold proteins involved in functions operating across the cell [50], which is why these were categorized as pleiotropic mediators of protein-protein interactions. Finally, as expected, only few genes encoding a metabolic function are upregulated in the three independent quiescent samples, including LINF_020008400, LINF_140012700, and LINF_350044100 encoding a drug/metabolite transporter, fatty acid elongase, and nucleoside diphosphate kinase, respectively. The upregulation of a fatty acid elongase has been previously described for quiescent parasites in murine lesions [36] and may be important for maintaining a favorable parasite/host interaction as *Leishmania* polyunsaturated fatty acid metabolites are important for macrophage M2 polarization. The functional annotation of an upregulated drug/metabolite transmembrane transporter that may be implicated in conferring quiescent parasites a higher tolerance to antileishmanial treatment. We propose that these upregulated genes participate in entry or maintenance of a quiescent state and may serve as positive markers of quiescence. Both positive and negative transcriptional markers or candidate drivers of entry and/or maintenance of quiescence warrant further investigation using the genetic toolbox available for *Leishmania* and may offer unprecedented insights in the universal problem of quiescence across the microbiological spectrum.

## METHODS

### Ethical statement

The use of laboratory rodents was carried out in strict accordance with all mandatory guidelines (EU directives, including the Revised Directive 2010/63/EU on the Protection of Animals used for Scientific Purposes that came into force on 01/01/2013, and the declaration of Helsinki in its latest version) and was approved by the Ethical Committee of the University of Antwerp, Belgium (UA-ECD 2019–04). Human bone marrow aspirate rest samples, obtained as a diagnostic sample without a written informed consent, were available for *in vitro* infection experiments following approval by the Committee of Medical Ethics UZA-UA (B3002021000027).

### Leishmania parasites

The *L. infantum* strains MHOM/FR/96/LEM3323 and MHOM/ES/2016/LLM2346 were kindly provided respectively by CNRL (Montpellier, France) and by WHOCC (Madrid, Spain), the latter being a recent clinical isolate. The *L. donovani* strain MHOM/ET/67/L82 was isolated from an Ethiopian VL patient. All were modified to express bioluminescent (PpyRE9) and fluorescent (DsRed) reporter genes integrated into the 18S rDNA locus (LEM3323 WT^PpyRE9/DsRed^, LLM2346 WT^PpyRE9/DsRed^ and Ldl82 WT^PpyRE9/DsRed^) [51, 52]. Promastigotes were sub-cultured twice weekly at 25°C in hemoflagellate-modified minimal essential medium (HOMEM, Gibco), supplemented with 10 % inactivated fetal calf serum (iFCS), 200 mM L-glutamine, 16.5 mM NaHCO_3_, 40 mg/L adenine, 3 mg/L folic acid, 2 mg/L D-biotin and 2.5 mg/L hemin. The number of passages was kept as low as possible to maintain parasite virulence.

### Laboratory animals and sand fly colony

Female BALB/c mice (6-8 weeks old) were purchased from Janvier (Genest-Saint-Isle, France) and accommodated in individually ventilated cages in SPF conditions. They were provided with food for laboratory rodents (Carfil, Arendonk, Belgium) and water *ad libitum*. Animals were subdivided in experimental groups based on simple randomization. Mice were kept in quarantine for at least 5 days before starting the experiment. Euthanasia was performed in CO_2_ chambers followed by cervical dislocation, and tissues were collected under aseptic conditions.

A *Lutzomyia longipalpis* sand fly colony was initiated with the kind help of NIH-NIAID (Prof. Shaden Kamhawi and Prof. Jesus Valenzuela) and maintained at the University of Antwerp under standard conditions (26°C, >75% humidity, in the dark) with provision of a 30% glucose solution *ad libitum* [53]. For infection experiments, 3-to 5-day old females from generations 31 to 44 were used.

### Primary mouse cells

Mouse BM was collected from BALB/c mice using two distinct techniques, based on pilot studies comparing alternative methods in terms of yield and quality. For both techniques, mice were sacrificed, and hind legs aseptically removed. Isolated femurs and tibias were cleaned by removing soft tissue from the bone using 70% ethanol-soaked cloth and tweezers.

For the <u>crushing</u> technique, the protocol was adapted from Lo Celso and Scadden [54]. Briefly, bones were crushed with mortar and pestle in ammonium-chloride-potassium (ACK) buffer (0.15 M NH_4_Cl, 1.0 mM KHCO_3_, 0.1 mM Na_2_EDTA) for erythrocyte lysis. Single cell suspensions were obtained by filtering through MACS® SmartStrainers (100 μm, Miltenyi Biotec), centrifuged at 500×*g* for 10 min (4°C) and resuspended in phosphate-buffered saline (PBS) + 0.2% bovine serum albumin (BSA). For efficient depletion of mature lineage-positive hematopoietic cells and to specifically isolate the preferred lineage-negative cells (*i.e.* undifferentiated progenitor cells), the Direct Lineage Cell Depletion Kit (Miltenyi Biotec) was employed according to manufacturer’s instructions. Following lineage depletion, cells were counted in PBS using a KOVA® counting chamber and resuspended in PBS + 0.2% BSA buffer to 2×10^7^ cells/mL. Cells were kept on ice during all procedures.

The <u>centrifugation</u> method, adjusted from the protocol described by Amend *et al.* [55] and Dobson *et al.* [56], was used for subsequent macrophage and dendritic cell differentiation. Briefly, a 0.5 mL microcentrifuge tube was perforated at the bottom with a 21G needle and nested inside a 1.5 mL tube (both from Eppendorf). After collection of femurs and tibias, one proximal end (knee epiphysis) was cut-off and placed in the 0.5 mL tube. Nested tubes were centrifuged in a microcentrifuge at 10,000×*g* for 15 sec, resulting in a visible pellet in the 1.5 mL tube. This pellet was then resuspended in ACK buffer for erythrocyte lysis.

To obtain BM-derived macrophages (BMDM), cells were centrifuged at 500×*g* for 10 min at 4°C, resuspended in Roswell Park Memorial Institute (RPMI) medium (Gibco) and divided over Petri dishes (Starstedt) supplemented with BM medium [RPMI 1640 medium with 10% (v/v) iFCS, 1% non-essential amino acids (NEAA), 1% sodium pyruvate, 1% L-glutamine, 50 U/mL penicillin, 50 μg/mL streptomycin (all from Gibco) and 15% L929 supernatant with M-CSF]. Following a 6-day incubation at 37°C with 5% CO_2_, the macrophages were collected by replacing the BM medium with ice cold dissociation buffer [PBS with 1% 0.5 M ethylenediaminetetraacetic acid (EDTA) and 2% 1 M 4-(2-hydroxyethyl)-1-piperazine-ethanesulfonic acid (HEPES)]. After detachment, the macrophage cell suspension was centrifuged at 500×*g* for 10 min and resuspended in RPMI medium. Cells were seeded in a 96-well plate (3×10^4^ cells/well) or a 24-well plate (1×10^6^ cells/well) and incubated for 24 h at 37°C with 5% CO_2_ to allow adherence of the BMDMs.

Primary peritoneal macrophages were obtained from Swiss mice after inoculation of 1 mL 2% starch solution in PBS. Macrophages were seeded in a 96-well plate (6×10^4^ cells/well) and kept at 37°C and 5% CO_2_ to allow adhesion. After 48 hours, macrophages were infected as described below.

### Primary human BM cells

Human BM aspirate was obtained from the iliac crest using BD Vacutainer® Plastic K3EDTA Tubes, initially collected for diagnostics, and delivered as residual sample. The BM was subjected to erythrocyte lysis twice using ACK buffer. Single cell suspensions were obtained by filtering through MACS® SmartStrainers (100 μm, Miltenyi Biotec), centrifuged at 300×*g* for 10 min (4°C) and resuspended in PBS + 0.2% BSA. Cells were counted in PBS and diluted to 2×10^7^ cells/mL for flow cytometric analysis. Cells were kept on ice during these procedures.

### In vitro and in vivo Leishmania infections

Parasite density was assessed using a KOVA® counting chamber. For *in vitro* infections, macrophages, LT-HSC and human hematopoietic stem and progenitor cells (HSPC) were co-cultured with stationary-phase promastigotes at a multiplicity of infection (MOI) of 5 for a minimum of 24h at 37°C with 5% CO_2_. For *in vivo* infection, stationary-phase parasites were centrifuged for 10 min at 4,000×*g* (25°C) and resuspended to 1×10^9^ parasites/mL in sterile RPMI medium. Mice were infected intravenously (i.v.) in the lateral tail vein with 1×10^8^ parasites in 100 µL of RPMI medium. Animals were monitored using *in vivo* bioluminescence imaging (BLI) at selected time points. Imaging was performed 3 min after intraperitoneal (i.p.) injection of 150 mg/kg D-Luciferin (Beetle Luciferin Potassium Salt, Promega) in the IVIS® Spectrum In Vivo Imaging System under 2.5% isoflurane inhalation anesthesia using 15 min exposure. Images were analysed using LivingImage v4.3.1 software by drawing regions of interests (ROIs) around specific organs to quantify the luminescent signal as relative luminescence units (RLU).

### Cell staining, flow cytometry and fluorescence-activated cell sorting (FACS)

Parasite cultures were analysed on a MACSQuant® Analyzer 10 (Miltenyi Biotec) after a 10 min centrifugation at 4,105×*g* and resuspension in PBS + 0.2% BSA buffer. Analyses were performed using FlowLogic^TM^ Software (Miltenyi Biotec) using a specific gating for singlet parasites expressing dsRed, for which the non-transfected parental parasite line served as a control. In some experiments, parasites were stained with 5-(and 6)-carboxyfluorescein diacetate succinimidyl ester (CFSE; Cell Division Tracker Kit, BioLegend) according to manufacturer’s instructions. Briefly, lyophilized CFSE was reconstituted with DMSO to a stock concentration of 5 mM. This stock solution was diluted in PBS to a 5 µM working solution. Promastigotes at a concentration of 10^8^ cells/mL were centrifuged at 4,000×*g* for 10 min and resuspended in CFSE working solution for 20 min at 25°C. The staining was quenched by adding 5 times the original staining volume of cell culture medium containing 10% FBS. Parasites were centrifuged again and resuspended in pre-warmed HOMEM medium for 10 min. After incubation, CFSE labeled parasites were used for infection and determining *in situ* proliferation in macrophages, LT-HSC and human HSPC.

Sorting of mouse LT-HSC and human HSPC was performed, and quality confirmed as described previously [16]. Briefly, BM cell suspensions (2×10^7^/mL concentration) were treated with FcɣR-blocking agent (anti-CD16/32, clone 2.4G2, BD Biosciences) for 15 min, followed by a washing step using 400×*g* centrifugation and resuspension in PBS + 0.2% BSA buffer. Next, cells were incubated for 20 min at 4°C with a mix of fluorescent conjugated anti-mouse antibodies at optimized concentrations. DAPI Staining Solution (Miltenyi Biotec) was used to assess viability. Cells were sorted using FACSMelody^TM^ (BD Bioscience) following specific gating strategies, confirmed with fluorescence minus one (FMO) controls and compensated using single stains. For visualizing infection, LT-HSC were collected on slides by Cytospin^TM^, fixed using methanol and stained for 15 min with Giemsa (Sigma Aldrich). Microscopic images were acquired using the UltraVIEW VoX dual spinning disk confocal system (PerkinElmer). For analysis of dsRed and/or CFSE levels on amastigotes at designated timepoints, infected macrophages and LT-HSC were recovered from the cultures. Cells were centrifuged at 400×*g* for 10 min and in PBS + 0.2% BSA. Host cell membranes were disrupted by 3 passages through a 25G needle. Amastigotes were collected in the supernatant after centrifugation at 250×*g* for 10 min and subsequently pelleted at 3,000×*g* for 10 min and resuspended in 500 µL PBS + 0.2% BSA for analysis by flow cytometry.

### RNA isolation and sequencing

Total RNA was extracted from three independent samples of 10,000 sorted DsRed^hi^ amastigotes or quiescent amastigotes (*L. infantum*) obtained from 5,000 sorted and infected LT-HSC. Extraction was performed with the QIAamp® RNA Blood Mini kit (Qiagen), according to the manufacturer’s instructions. To exclude gDNA, an additional step using gDNA elimination columns (Monarch®) was performed. RNA samples were stored in aliquots at –80°C. Unbiased total RNA sequencing was performed at Brightcore using the SMARTer® Stranded RNA-Seq Kit to generate strand-specific RNAseq libraries for Illumina® sequencing. Reads were generated in an S4 run (2×100bp, 200M reads) on an Illumina NovaSeq 6000 apparatus. Alignments of the low input RNAseqs with the *L. infantum* JPCM5 reference genome were made using BWA (Burrows-Wheeler Aligner) and normalization was performed in DESeq2 using the variance stabilizing transformation (VST) in the default unsupervised mode [57]. Differential expression analysis was largely based on a workflow using Bioconductor packages in R [58]. Euclidean distance between samples was calculated using the R function *dist* and with the Poisson Distance package *PoiClaClu* (https://CRAN.R-project.org/package=PoiClaClu) and visualized in a heatmap using *pheatmap* (https://CRAN.R-project.org/package=pheatmap) and the colorRampPalette function from the RColorBrewer (https://CRAN.Rproject.org/package=RColorBrewer). PCA and Volcano plots were generated using *ggplot2* (https://CRAN.Rproject.org/package=ggplot2) and *ggrepel* (https://cran.r-project.org/web/packages/ggrepel/). Heatmaps of the top 150 differential genes and the top 25 upregulated and 25 downregulated genes were generated using *pheatmap*. Gene Set Enrichment by Ontology Analysis was performed in R, using the topGO library, based on Fisher’s exact test. The false discovery rate correction was performed using Benjamini-Hochberg, and the p.adjust function, in R.

### Functional annotation through bio-informatics and structure prediction

Insights in differentially expressed genes were obtained by searches for functional annotation or orthologues in TriTrypDB [59]. An approach combining bioinformatics and AlphaFold2 [24, 25] structure prediction, followed by a structural homology search using the DALI server [60] was employed to elucidate the putative roles of the proteins encoded by upregulated quiescence-associated genes without annotated function. Amino acid sequences were subjected to a search on the InterPro web server (https://www.ebi.ac.uk/interpro/) to find potential functional motifs. Protein structures for all 24 candidates were predicted using Alphafold2 and their (local) quality were critically assessed based on the pLDDT confidence metric (score between 0 and 100 that estimates how well predicted models agree with experimentally determined structures). AlphaFold-Multimer [61] was employed for one candidate (LINF_230021900) to generate a structural model of a homodimer and dimer formation was critically assessed based on the predicted aligned errors (PAE) confidence metric. The structural models were then submitted to the DALI web server for a structural homology search and hits were carefully evaluated based on the DALI Z-score and Cα rmsd. Molecular graphics and analyses were performed with UCSF ChimeraX [62, 63].

### Sand fly infections and evaluation of parasite load

Sand fly females (*L. longipalpis*) were fed with heat-inactivated heparinized mouse blood containing 5 × 10^6^/mL promastigotes from log-phase cultures through a chicken skin membrane. Groups were randomized by an independent researcher until data analysis to avoid bias. Blood-fed females were separated 24 h after feeding, kept in the same conditions as the colony and dissected on 5, 7, 9, and 12 days post blood meal to microscopically check the presence of parasites. Following disruption of the total gut in 50 μL PBS, the parasite load was quantified microscopically using a KOVA® counting chamber [64, 65]. Parasites isolated from sand flies on day 12 post blood meal were cultured in HOMEM promastigote medium supplemented with 5% penicillin-streptomycin, to obtain post sand fly cultures.

### Promastigote growth

Promastigote growth curves were made as described before [66] to compare the *in vitro* growth of quiescent *vs* non-quiescent strains. After passage through fine needles (21G and 25G) to break clustering, the promastigotes were diluted in PBS and counted by KOVA® counting chamber. Exactly 5×10^5^ log-phase promastigotes/mL were seeded in 5 mL HOMEM and their number was determined by microscopic counting every 24 h for a total of 10 days. Three independent repeats of each strain were run in parallel.

### Statistics and Reproducibility

Statistical analyses were performed using GraphPad® Prism version 9.0.1. Tests were considered statistically significant if *p* <0.05. Growth curves were statistically compared using Wilcoxon matched-pairs signed rank test. Parasite load in sand fly infections were tested using Unpaired *t* test. Infectivity in macrophages was tested using Ordinary one-way ANOVA. MFI of DsRed expression was compared using Mann-Whitney test.

## Data availability

The authors declare that the data underlying the findings of this study are available within the paper and its Supplementary Information files are available upon request. The source data underlying Figs. 1-6 and Supplementary Figs. 1-3 will be provided as Source Data file.

## ACKNOWLEDGEMENTS

This work was supported by the Fonds Wetenschappelijk Onderzoek (www.fwo.be; grant numbers 1S30721N and G065421N) and the University of Antwerp (www.uantwerpen.be; grant number TT-ZAPBOF 33049). J.C. and D.J. were funded by an MRC New Investigator Research Grant to DJ (MR/T016019/1). The authors would like to thank Pim-Bart Feijens for excellent technical assistance. LMPH is a partner of the Excellence Centre ‘Infla-Med’ (www.uantwerpen.be/infla-med).

## AUTHOR CONTRIBUTIONS

G.C. and L.M. conceived and supervised the study. G.C. and L.D. designed experiments, L.D., S.V.A and Y.N. performed experiments. S.H. completed sand fly infection studies. G.C., L.M. and D.E. contributed reagents. L.D., G.C, J.C., R.A., Y.S. and H.I. analyzed the data. G.C., L.M., D.E. and D.J. acquired project funding. B.C., H.I., J.C., D.J. participated in RNA sequencing experiments and analyses. Y.S. performed protein prediction studies. L.D., L.M., and G.C. prepared the manuscript.

## SUPPLEMENTAL INFORMATION TITLES AND LEGENDS

**Supplementary figure 1. Quiescent promastigotes lose DsRed marker. (a)** sorted human HSPC were infected for 24 hours with *L. infantum* (LLM1246 WT^PpyRE9/DsRed^). Amastigotes were recovered and measured via flow cytometry. **(b)** RT-qPCR on RNA samples extracted from recovered cultures after promastigote back-transformation from (a).

**Supplementary figure 2. Promastigote growth curves remain stable between quiescent and DsRed^hi^ strains. (a)** *In vitro* growth curves of LEM3323 promastigotes recovered from quiescent and non-quiescent (DsRed^hi^) parasites in infected mouse LT-HSC, both before (pre) and after (post) sand fly passage. **(b)** LLM2346 promastigote growth curves *in vitro* of quiescent and non-quiescent (DsRed^hi^) strains recovered from infected human HSPC. Wilcoxon matched-pairs signed rank test, \*\**p* <0.01. **(c)** LEM3323 promastigote growth curves *in vitro* of relapse and infection strains recovered from relapsed and infected BALB/c mice as described above. All results are based on three independent replicates.

**Supplementary figure 3. GO term analysis of quiescent Leishmania amastigotes.** Visual representation of GO terms enriched in biological processes **(a)** cellular components **(b)** and molecular function **(c)**. Interesting changes are highlighted in yellow.

